# Crystal structure of the LRR ectodomain of the plant immune receptor kinase SOBIR1

**DOI:** 10.1101/581231

**Authors:** Ulrich Hohmann, Michael Hothorn

## Abstract

Plant unique membrane receptor kinases with leucine-rich repeat (LRR) extracellular domains are key regulators of development and immune responses. Here we present the 1.55 Å resolution crystal structure of the immune receptor kinase SOBIR1 from Arabidopsis. The ectodomain structure reveals the presence of 5 LRRs sandwiched between non-canonical capping domains. The disulphide bond-stabilized N-terminal cap harbors an unusual β-hairpin structure. The C-terminal cap features a highly positively charged linear motif which we find largely disordered in our structure. Size-exclusion chromatography and right-angle light scattering experiments suggest that SOBIR1 is a monomer in solution. The protruding β-hairpin, a set of highly conserved basic residues at the inner surface of the SOBIR LRR domain and the presence of a genetic missense allele in LRR2, together suggest that the SOBIR1 ectodomain may mediate protein – protein interaction in plant immune signalling.

**Synopsis:** The ectodomain structure of a novel plant membrane receptor kinase with unusual capping domains is reported.

## 1. Introduction

Plants have evolved a unique set of membrane receptor kinases (LRR-RKs) that are composed of a leucine-rich repeat ectodomain, a transmembrane helix and a dual-specificity kinase domain in the cytoplasm (Shiu & Bleecker, 2001). The ectodomains of LRR-RKs show a bimodal size distribution (Fig. 1*a*). Family members with large ectodomains (15-30 LRRs) represent ligand binding receptors (Hohmann *et al.*, 2017). In contrast, SOMATIC EMBRYOGENESIS RECEPTOR KINASES (SERKs) (Schmidt *et al.*, 1997) with short ectodomains (5 LRRs) have been characterized as essential co-receptors (Brandt & Hothorn, 2016). Ligand binding to large LRR-RKs promotes their association with shape-complementary SERKs at the cell surface, which in turn enables their cytoplasmic kinase domains to interact and to trans-phosphorylate each other (Santiago *et al.*, 2013, 2016; Hohmann, Santiago *et al.*, 2018). SERKs represent only five of the ∼60 small LRR-RKs in Arabidopsis (Fig. 1*a*), but genetic evidence suggests that sequence-related NIK/CIK/CLERK proteins may fulfil similar functions (Hu *et al.*, 2018; Cui *et al.*, 2018; Anne *et al.*, 2018).

**Figure 1.**
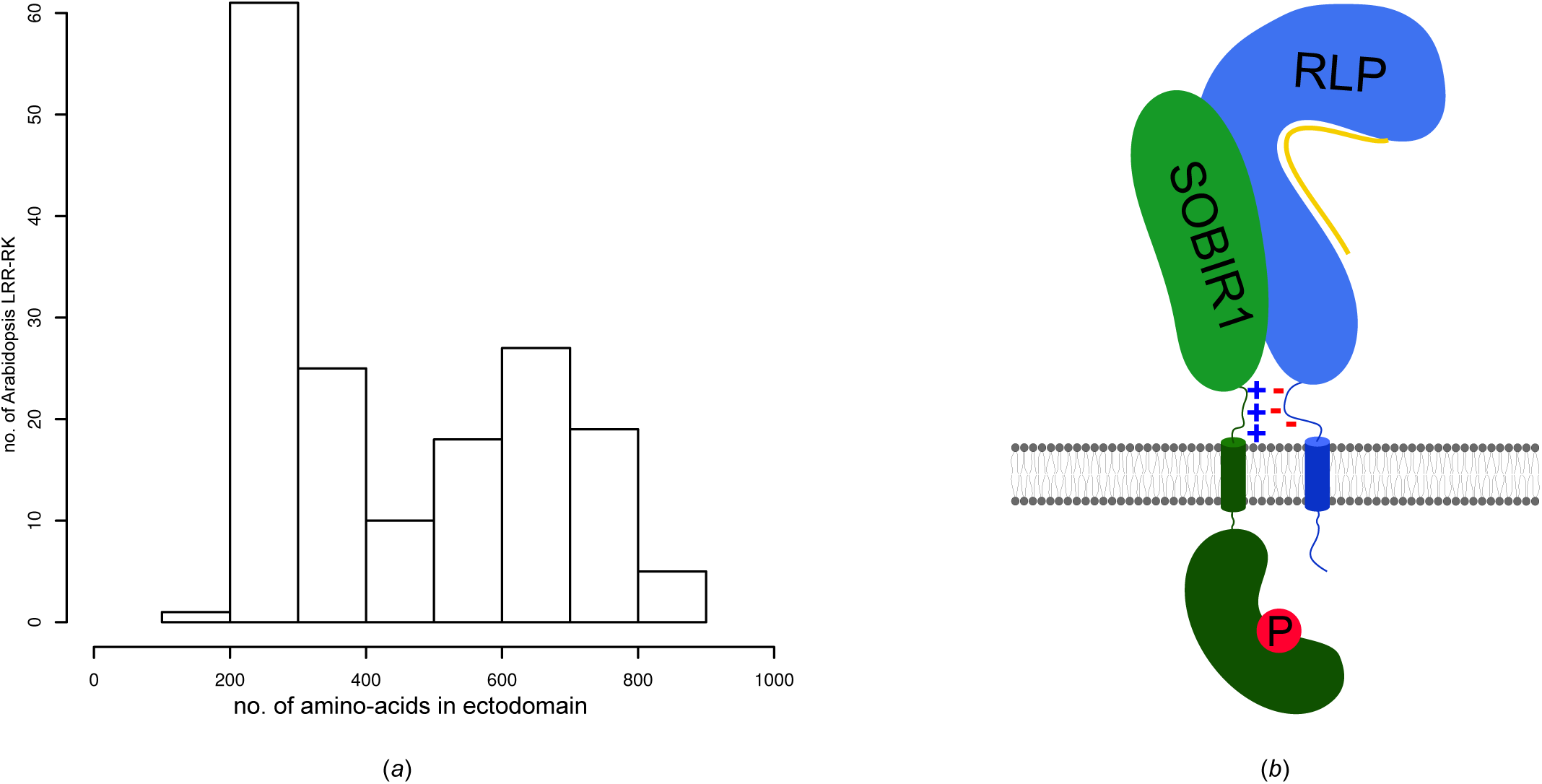
Distribution of leucine-rich repeat receptor kinases in Arabidopsis. (*a*) Histogram showing the distribution of Arabidopsis LRR-RKs by ectodomain size. Plotted are the the number of residues in the ectodomain vs. the number of LRR-RKs found in the current reference proteome of *Arabiodpsis thaliana*. (*b*) Cartoon model for a putative SOBIR1 – receptor-like protein (RLP) – ligand (yellow) complex at the plasma membrane (shown in grey). SOBIR kinase domain (green, kidney-shaped) phosphorylation is indicated with a P. Note the presence of opposing charged stretches (+ and -) next to the transmembrane helices (cylinders) in SOBIR1 and different RLPs.

Recently, the BIR family of receptor pseudo-kinases (5 LRRs in the ectodomain), have been defined as negative regulators of SERK co-receptors (Ma *et al.*, 2017; Hohmann, Nicolet *et al.*, 2018). Ligand-independent interaction of a BIR and a SERK ectodomain keeps the LRR-RK co-receptor in a basal, inhibited state (Hohmann, Nicolet *et al.*, 2018). In addition, the structure of POLLEN RECEPTOR-LIKE KINASE 6 (6 LRRs in the ectodomain) in complex with a peptide hormone ligand has been reported, but it is presently unclear if PRK6 represents the receptor or a co-receptor for these peptides (Zhang *et al.*, 2017).

Here we report the structure of the functionally distinct plant receptor kinase SOBIR1, predicted to have 4 LRRs in its ectodomain (Gao *et al.*, 2009). SOBIR1 was initially found in a suppressor screen of the *bir1-1* mutant, which displays autoimmune phenotypes (Gao *et al.*, 2009). SOBIR1 loss-of-function restored wild-type like growth in *bir1-1*, suggesting that SOBIR1 functions in plant immune signalling (Gao *et al.*, 2009). Subsequently, it was found that SOBIR1 interacts with receptor-like proteins (RLPs) (Liebrand *et al.*, 2013), a family of plant membrane proteins (∼ 50 family members in Arabidopsis) that harbour LRR ectodomains and a trans-membrane helix, but which lack a cytoplasmic kinase domain (Wang *et al.*, 2008; Gust & Felix, 2014) (Fig. 1*b*). Many RLPs are plant immune receptors that recognize various microbe-associated molecular patterns and whose signalling function depends on SOBIR1 (Zhang *et al.*, 2013; Jehle *et al.*, 2013; Zhang *et al.*, 2013; Albert *et al.*, 2015; Catanzariti *et al.*, 2017; Wang *et al.*, 2018; Domazakis *et al.*, 2018). How different RLPs interact with SOBIR1 to activate plant immune signalling is poorly understood at the molecular level. Presently, it is known that a GxxxG motif in the SOBIR1 trans-membrane helix is required for the interaction with different RLPs (Bi *et al.*, 2016). It has also been demonstrated that the kinase activity of SOBIR1 is essential for its signalling function (van der Burgh *et al.*, 2019). Here, we present the crystal structure of the SOBIR1 ectodomain from *Arabidopsis thaliana* and discuss its implications for plant immune signalling.

## 2. Material and methods

### 2.1 Analysis of LRR ectodomain size-distribution

166 *Arabidopsis thaliana* proteins containing predicted N-terminal LRR and C-terminal kinase domains connected via a single trans-membrane helix were identified in Araport11 (http://arabidopis.org). The LRR ectodomain sequences were isolated by defining putative signal peptides using SignalP (http://www.cbs.dtu.dk/services/SignalP/, version. 5.0) (Armenteros *et al.*, 2019) and transmembrane helices using TMHMM (http://www.cbs.dtu.dk/services/TMHMM/, version 2.0) (Möller *et al.*, 2001). Data were plotted in R (R Core Team, 2014) (Fig. 1*a*).

### 2.2 Protein expression and purification

The coding sequences of AtSOBIR1 (residues 1-270 and 1-183) as well as AtRLP23^1-849^ and AtRLP32^1-818^ were amplified from *Arabidopsis thaliana* genomic DNA, gp67-RLP23^23-847^, codon optimized for expression in *Spodoptera frugiperda*, was obtained from T. Nürnberger and PpNLP20-stop, codon optimized for expression in *Spodoptera frugiperda*, was obtained as a synthetic gene (Twist Bioscience, San Francisco, USA). All protein coding sequences were cloned into a modified pFastBac vector (Geneva Biotech), which provides a *tobacco etch virus* protease (TEV)-cleavable C-terminal StrepII-9xHis tag. Shortened expression constructs and signal peptide swaps were constructed using Gibson-assembly cloning strategies (Gibson *et al.*, 2009).

For protein expression, *Trichoplusia ni* (strain Tnao38) (Hashimoto *et al.*, 2010) cells were infected with 15 ml of virus in 250 ml of cells at a density of 2.3 × 10^6^ cells ml^−1^, incubated for 26 h at 28 °C and 110 rev min^−1^ and then for another 48 h at 22 °C and 110 rev min^−1^. For co-expression, cells were infected with 10 ml of each virus. Subsequently, the secreted ectodomains were purified from the supernatant by sequential Ni^2+^ (HisTrap excel; GE Healthcare; equilibrated in 25 mM KP_i_ pH 7.8, 500 mM NaCl) and StrepII (Strep-Tactin XT; IBA; equilibrated in 25 mM Tris pH 8.0, 250 mM NaCl, 1 mM EDTA) affinity chromatography. The proteins were further purified by size-exclusion chromatography (on either a Superdex 200 increase 10/300 GL or a HiLoad 16/600 Superdex 200pg column, both GE Healthcare), equilibrated in 20 mM sodium citrate pH 5.0, 150 mM NaCl. Purified proteins were then concentrated using Amicon Ultra concentrators (molecular-weight cutoff 10,000; Millipore) and purity and structural integrity was assessed by SDS-PAGE and Right Angle Light Scattering (RALS). The molecular weight of the proteins (as determined by RALS) is ∼ 30.2 kDa for AtSOBIR^1-270^ and ∼ 38.1 kDa for AtSOBIR^1-283^.

### 2.3 Crystallisation and crystallographic data collection

Hexagonal SOBIR1 crystals (∼400 × 80 × 80 μm) developed in hanging drops composed of 1 μl protein solution (20 mg ml^−1^ in 20 mM sodium citrate pH 5.0, 150 mM NaCl) and 1 μl crystallisation buffer (25 % [w/v] PEG 3,350, 1M LiCl, 0.1 M sodium acetate pH 5.5) suspended of 1 ml of the latter as reservoir solution. Crystals were cryoprotected by serial transfer in crystallisation buffer supplemented with glycerol to a final concentration of 15 % (v/v), and snap-frozen in liquid N_2_.

Native (λ=1.033201 Å, 1 360° wedge at 0.1° oscillation) and redundant sulphur single-wavelength anomalous dispersion (SAD) data (λ= 2.078524 Å, 3 360° wedges at 0.1° oscillation) to 1.75 Å and 3.12 Å resolution, respectively were collected at beam line X06DA of the Swiss Light Source (SLS), Villigen, Switzerland equipped with a Pilatus 2M-F detector (Dectris Ltd.) and a multi-axis goniometer. A second native dataset at 1.55 Å resolution was recorded from a different crystal. Data processing and scaling was done with XDS and XSCALE, respectively (version January, 2018) (Kabsch, 1993).

### 2.4 Structure determination and refinement

The anomalous signal in the scaled SAD dataset extended to ∼ 4.0 Å resolution when analysed with phenix.xtriage (Zwart *et al.*, 2005; Adams *et al.*, 2010). The structure was solved using the molecular replacement (MR)-SAD method as implemented in Phaser (McCoy *et al.*, 2007). An alignment of SOBIR1 and SERK1 ectodomains (sharing ∼30 % sequence identity) was prepared in HHPRED (Zimmermann *et al.*, 2018) and input into the program CHAINSAW (Stein, 2008). The LRR ectodomain of SERK1 (PDB-ID 4lsc) with non-identical side-chains trimmed to alanine, was used as search model. Phaser returned a single solution in space-group *P*6_5_, comprising a dimer in the asymmetric unit. The first molecule had a Phaser rotation function Z-score (RFZ) of 3.5, a translation function Z-score (TFZ) of 6.6 and an associated log-likelihood gain (LLG) of 60. The TFZ for the second molecule was 14.4 with a final refined LLG of 351. The resulting partial model (starting figure of merit [FOM] was 0.24 at 3.12 Å resolution) was used to locate nine putative sulphur sites by log-likelihood-gradient completion in Phaser (final FOM was 0.46). Density modification and phase extension to 1.75 Å resolution in phenix.resolve (Terwilliger, 2003) yielded a readily interpretable electron density map. The structure was completed by alternating cycles of manual model building and correction in COOT (Emsley & Cowtan, 2004) and restrained TLS refinement against a 1.55 Å resolution native dataset in phenix.refine (Afonine *et al.*, 2012) (Table 1). Inspection of the final model with phenix.molprobity (Davis *et al.*, 2007) revealed excellent stereochemistry (Table 1). Structural representations were done in Pymol (http://pymol.org) and Chimera (Pettersen *et al.*, 2004). Electrostatic potentials were calculated using the Pymol APBS plugin (Jurrus *et al.*, 2018). To visualize phased anomalous difference maps, |*F*_A_| values and phase shifts were calculated from the SAD dataset in XPREP (Bruker) and together with the final SOBIR1 coordinate file input into the program AnoDe (Thorn & Sheldrick, 2011). The resulting map file was converted to ccp4 format using the shelx2map.

**Table 1.** Crystallographic data collection, phasing and refinement statistics

|  | Sulphur SAD | Native 1 | Native 2 (high-res) |
| --- | --- | --- | --- |
| **Data collection** |  |  |  |
| Space group | | $P6_5$ | $P6_5$ |
| Cell dimensions |  |  |  |
| a, b, c (Å) | 81.9, 81.9, 109.8 | 81.9, 81.9, 109.8 | 82.3, 82.3, 109.9 |
| $\alpha, \beta, \gamma$ (°) | 90, 90, 120 | 90, 90, 120 | 90, 90, 120 |
| Resolution (Å) | 43.41 – 3.12 (3.18 – 3.12) | 41.43 – 1.75 (1.80 – 1.75) | 43.5 – 1.55 (1.65 – 1.55) |
| $R_{\text{meas}}^{\#}$ | 0.07 (0.23) | 0.14 (3.59) | 0.07 (2.73) |
| CC(1/2)<sup>#</sup> (%) | 100 (100) | 100 (64.9) | 100 (51.3) |
| $I/\sigma I^{\#}$ | 47.05 (13.4) | 17.6 (1.0) | 25.0 (0.9) |
| Completeness (%)<sup>#</sup> | 100 (100) | 100 (99.9) | 99.9 (99.3) |
| Redundancy<sup>#</sup> | 29.2 (22.6) | 21.7 (21.1) | 18.5 (12.6) |
| Wilson B-factor<sup>#</sup> | 37.2 | 38.0 | 34.1 |
| **Phasing** |  |  |  |
| Resolution (Å) | 43.41 – 3.12 |  |  |
| No. of sites | 9 |  |  |
| FOM<sup>*</sup> | 0.428 |  |  |
| **Refinement** |  |  |  |
| Resolution (Å) |  |  | 43.5 – 1.55 (1.58 – 1.55) |
| No. reflections |  |  | 60,266 (2,528) |
| $R_{\text{work}}/R_{\text{free}}^{\S}$ | | | 0.172/0.188 (0.416/0.423) |
| No. atoms |  |  |  |
| Protein |  |  | 2,949 |
| Glycan |  |  | 28 |
| Buffer |  |  | 15 |
| Chloride |  |  | 1 |
| Water |  |  | 254 |
| Res. B-factors<sup>\S</sup> |  |  |  |
| Protein |  |  | 40.2 |
| Glycan |  |  | 85.1 |
| Buffer |  |  | 69.1 |
| Chloride |  |  | 32.1 |
| Water |  |  | 43.1 |
| R.m.s deviations<sup>\S</sup> |  |  |  |
| Bond lengths (Å) |  |  | 0.005 |
| Bond angles (°) |  |  | 1.05 |
| Molprobity results |  |  |  |
| Ramachandran outliers (%) |  |  | 0 |
| Ramachandran favored (%) |  |  | 95.24 |
| Molprobity score |  |  | 1.40 |
| PDB code |  |  | **6r1h** |

<sup>#</sup>as defined in XDS (Kabsch, 1993)
<sup>\*</sup>as defined in Phaser (McCoy *et al.*, 2007)
<sup>\S</sup>as defined in phenix.refine (Afonine *et al.*, 2012)

### 2.5 Analytical size-exclusion chromatography

Analytical size exclusion chromatography (SEC) experiments were performed on a Superdex 200 increase 10/300 GL column (GE Healthcare) pre-equilibrated in 20 mM sodium citrate pH 5.0, 250 mM NaCl. 200 μg of protein, injected in a volume of 100 μl, was loaded onto the column and elution at 0.75 ml min^−1^ was monitored by ultraviolet absorbance at λ = 280 nm. Peak fractions were analysed by SDS-PAGE.

### 2.6 Right-angle light scattering

The oligomeric state of SOBIR1 was analysed by size exclusion chromatography paired with a right-angle light scattering (RALS) and a refractive index (RI) detector, using an OMNISEC RESOLVE / REVEAL combined system. Calibration of the instrument was carried out using a BSA standard (Thermo Scientific Albumin Standard). 100 μg of protein, in a volume of 50 μl, were separated on a Superdex 200 increase column (GE Healthcare) in 20 mM sodium citrate pH 5.0, 250 mM NaCl at a column temperature of 35 °C and a flow rate of 0.7 mL/min. Data was analysed using the OMNISEC software (v10.41).

## 3. Results

We obtained the *Arabidopsis thaliana* SOBIR1 ectodomain (residues 1-270) by secreted expression in insect cells (Fig. 2*a*, see Methods). The N-glycosylated protein was crystallized using the vapour diffusion method and the structure was solved by molecular replacement – single wavelength anomalous dispersion (MR-SAD) at beam line X06DA of the Swiss Light Source (see Methods, Table 1) (Basu *et al.*, 2019). The solution in space-group *P*6_5_ comprises a dimer in the asymmetric unit, with the nine putative sulphur sites corresponding to a disulphide-bridge in the N-terminal LRR capping domain, to a free cysteine and a methionine residue in the LRR core, and to a free ion, which we interpreted as a chlorine anion originating from the crystallisation buffer (Fig. 2*b*). The model was refined against an isomorphous, high-resolution native dataset at 1.55 Å resolution. An example region of the final (2*F*_o_ - *F*_c_) map is shown in Fig. 2*c*, highlighting the only N-glycan located in the structure, attached to asparagine 154.

**Figure 2.**
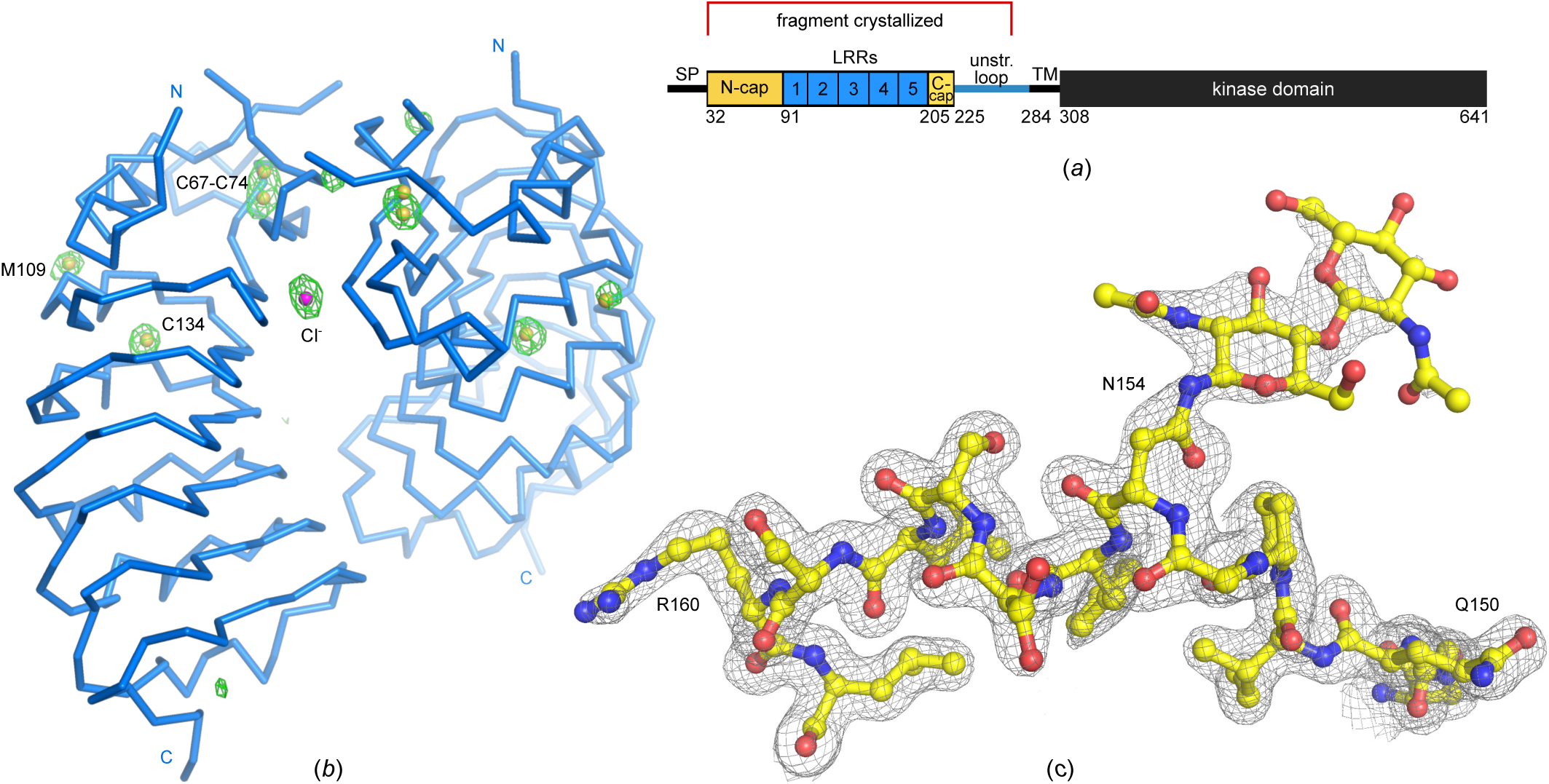
Structure solution of SOBIR1. (*a*) Schematic representation of AtSOBIR1 (SP, signal peptide; TM, transmembrane helix; unstr. loop, unstructured loop; N-cap / C-cap, N/C-terminal capping domain). The fragment crystallized is indicated in red. (*b*) C_α_ trace of the SOBIR1 crystallographic dimer (in blue) and including a phased anomalous difference map contoured at 5.0 σ (green mesh), the eight sulphur atoms (yellow spheres) and a putative chloride anion (in magenta). (*c*) Example region of the SOBIR1 structure, including the N-glycosylated Asn154, (yellow, in ball-and-stick representation) and including the final (2*F*_*o*_*-F*_*c*_) map contoured at 1.2 σ.

The refined model reveals the presence of 5 LRRs in the SOBIR1 ectodomain, not 4 as initially proposed (Gao *et al.*, 2009) (Fig. 3*a,b*). A genetic missense allele (*sobir1-8*, Val129 to Met), which causes a weak *sobir1* loss-of-function phenotype, maps the the outer face of the LRR core in LRR2 (Gao *et al.*, 2009). The SOBIR1 LRR core is masked by a N-terminal capping domain, as found in many plant LRR-RKs (residues 34-90, shown in yellow in Fig. 3*b*) (Hohmann *et al.*, 2017). Loop residues 57-63 appear disordered in our structure (shown in grey in Fig. 3*b*). The N-terminal cap features a protruding, unusual β-hairpin structure (shown in magenta in Fig. 3*b*), which presents several conserved basic and hydrophobic amino acids on its surface (Fig. 3*c*).

**Figure 3.**
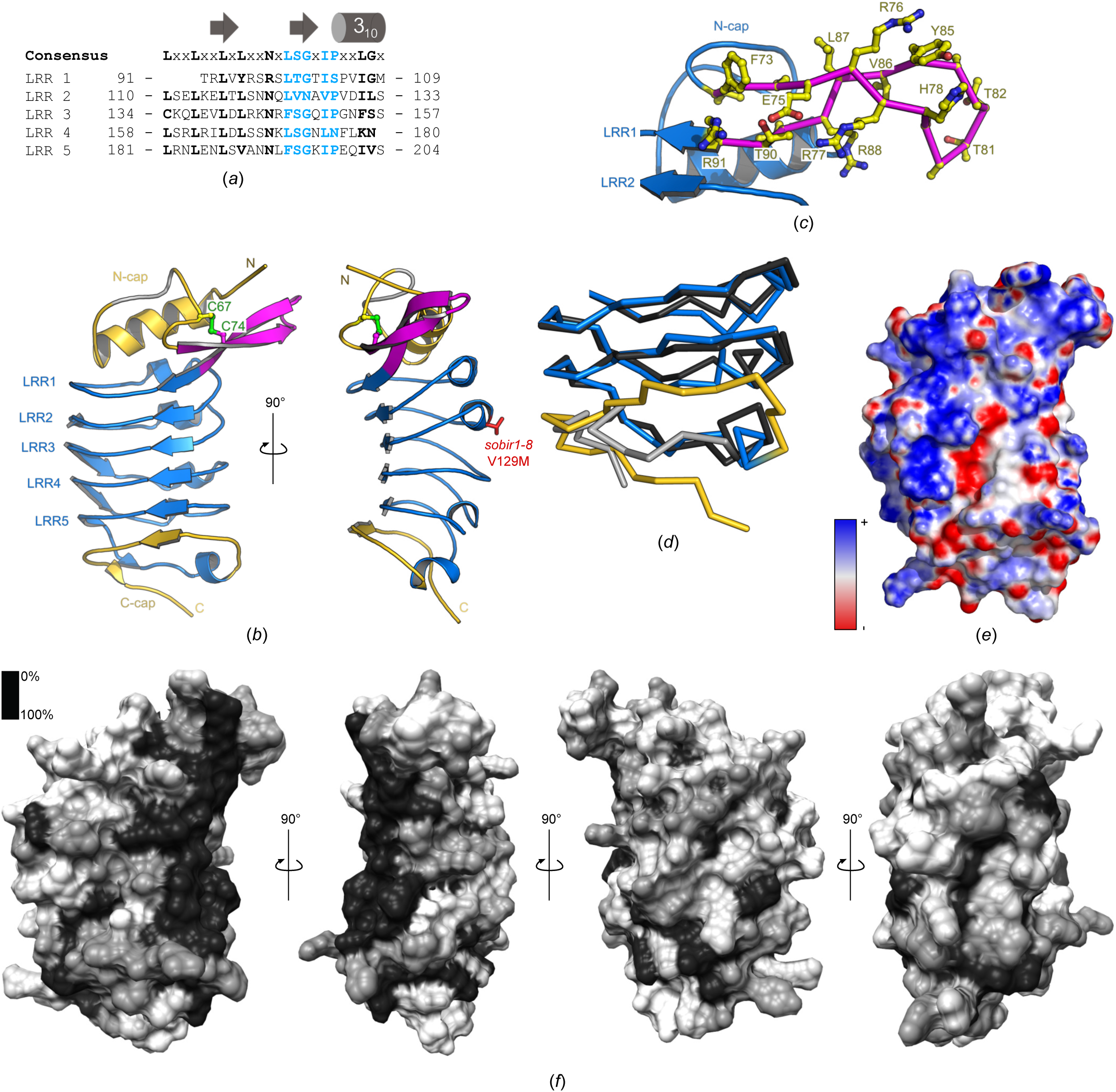
The SOBIR1 ectodomain harbours 5 LRRs and unusual capping domains. (*a*) Sequence alignment of the 5 SOBIR1 LRRs, with the canonical consensus sequence in black and the plant-specific LRR motif in blue. The LRR consensus sequence and a secondary structure assignment calculated with DSSP (Kabsch & Sander, 1983) are shown alongside. (*b*) Front (left) and y-axis rotated side (right) view of the SOBIR1 LRR domain (blue ribbon diagram), with N- and C-terminal capping domains shown in yellow, the SOBIR1-specific extended β-hairpin in magenta and a disordered loop in the N-terminal capping domain (residues 57-63) in grey. Shown in bond representation are the disulphide bond (in green) and Val129 (in red), which is mutated to Met in *sobir1-8*. (*c*) Close-up view of the extended β-hairpin (magenta, C_α_ trace, with side chains shown in ball-and-stick representation, in yellow) in the SOBIR1 ectodomain (blue ribbon diagram) (*d*) C_α_ traces of a structural superposition of the ectodomains of SOBIR1 (blue, C-terminal capping domain in yellow) and SERK3 (PDB-ID 4mn8, (Sun, Li *et al.*, 2013), black, C-terminal capping domain in grey). (*e*) Surface representation of the SOBIR1 ectodomain coloured according to the electrostatic surface potential (blue, negative; red, positive). (*f*) Surface representation of the SOBIR1 ectodomain coloured according to SOBIR1 sequence conservation, comparing SOBIR orthologues from different plant species (sequences in Fig. 4*c*). Note the presence of a highly conserved patch at the outer edge of the inner surface, ranging from the N-terminal cap through LRRs 1-5.

A highly basic, low complexity region is located at the C-terminus of SOBIR ectodomains from different plant species (Fig. 4c). In line with this, the C-terminal capping domain, which in plant LRR-RKs normally terminates by a well-defined disulphide bond (Hohmann *et al.*, 2017), is found largely disordered in our SOBIR1 structure (Fig. 3b). This is reminiscent of the LRR-RK SERK3, which contains a proline-rich sequence at the C-terminus of its ectodomain and whose C-terminal capping domain was also found largely unstructured in different SERK3 – LRR-RK complex structures (Fig. 3*d*) (Sun, Li *et al.*, 2013; Sun, Han *et al.*, 2013).

**Figure 4.**
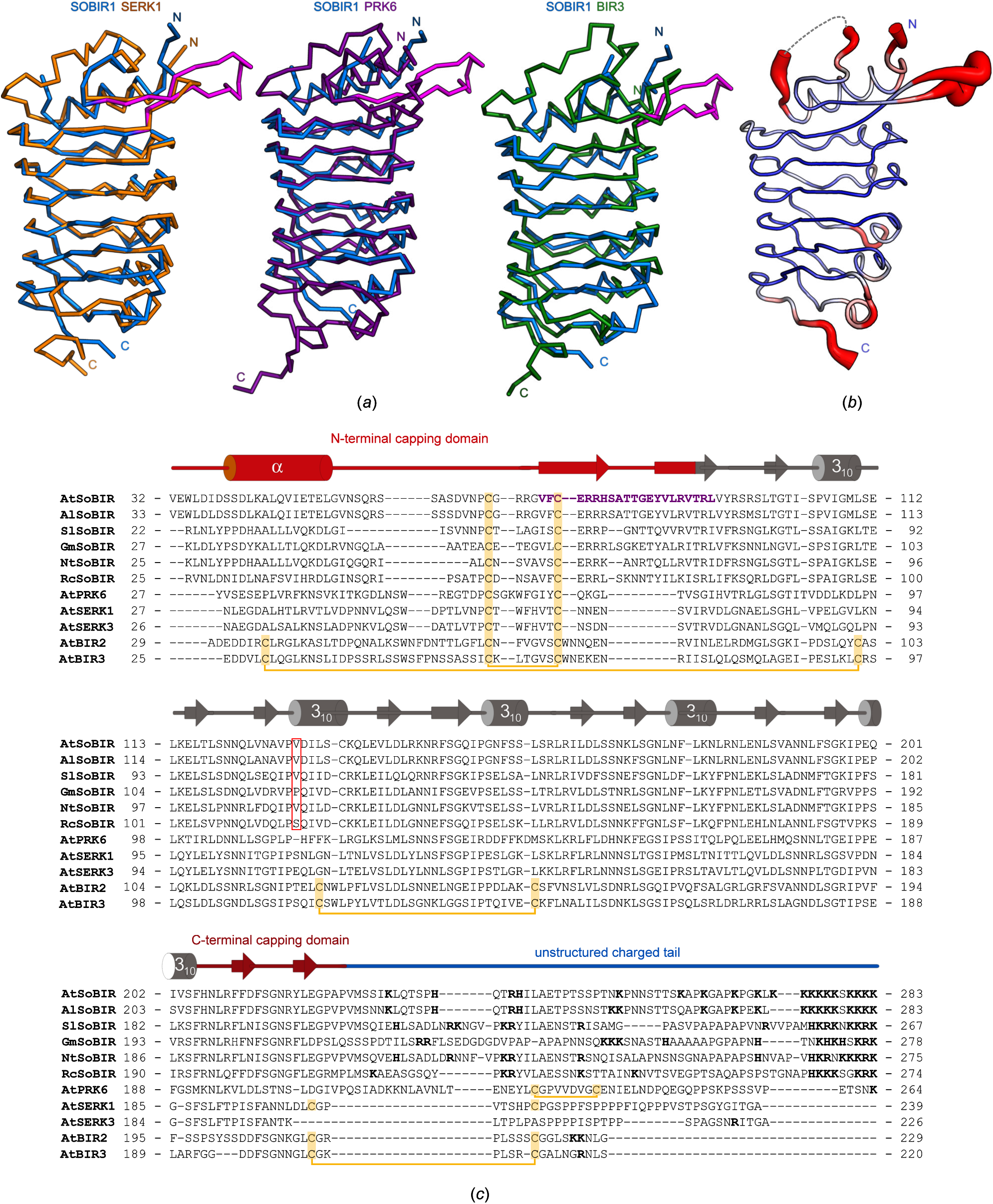
AtSOBIR1 shares a common architecture with other small plant LRR-RKs (*a*) C_α_ traces of structural superpositions of the ectodomain of SOBIR1 (blue) with AtSERK1 (left, in orange; PDB-ID 4lsc (Santiago *et al.*, 2013)), AtPRK6 (purple, middle; PDB-ID 5yah (Zhang *et al.*, 2017)) and AtBIR3 (green, right; PDB-ID 6fg8 (Hohmann, Nicolet *et al.*, 2018)). The SOBIR1-unique extended β-hairpin is highlighted in magenta. (*b*) Ribbon diagram of the SOBIR1 ectodomain with C_α_ atoms coloured according to their crystallographic temperature factors (red, high; blue, low). Residues 57-63, which are missing in the structure, are indicated by a dotted line. The N- and C-termini as well as a N-terminal capping domain loop and the extending β-hairpin appear flexible in contrast to the rigid and well-ordered LRR core. (*c*) Structure based sequence alignment of the ectodomains of SOBIR1 from *Arabidopsis thaliana* (Uniprot [http://www.uniprot.org] identifier: Q9SKB2), *Arabidopsis lyrata* (Uniprot identifier: D7LEA5), *Solanum lycopersicum* (Uniprot identifier: K4C8Q3), *Glycine max* (Uniprot identifier: I1JXE0), *Nicotiana tabacum* (Uniprot identifier: Q8LP72) and *Ricinus communis* (Uniprot identifier: B9RAQ8) as well as *Arabidopsis thaliana* SERK1 (Uniprot identifier: Q94AG2), SERK3 (Uniprot identifier: Q94F62), BIR2 (Uniprot identifier: Q9LSI9) and BIR3 (Uniprot identifier: O04567). Shown alongside is a secondary structure assignment (calculated with DSSP (Kabsch & Sander, 1983)), with the N- and C-terminal capping domains highlighted in red and a unstructured region at the C-terminus drawn in blue. Disulfide bridges are shown in yellow, the SOBIR1-specific β-hairpin in purple, the position of the AtSOBIR1^V129^, mutated to methionine in *sobir1-8*, is indicated by a red box; positively charged residues in the unstructured C-terminal tail are highlighted in bold and black.

Analysis of the electrostatic surface potential of the SOBIR1 ectodomain revealed several basic patches at the inner side of the LRR solenoid (Fig. 3*e*), some of which are highly conserved among SOBIR orthologs from different plant species (Fig. 3*f*,4*c*).

We next compared SOBIR1 to other plant LRR-RK ectodomains. A structural homology search with the program DALI (Holm & Sander, 1993) returned several large and small LRR ectodomains as top hits. We focused our analysis on plant LRR-RKs with small ectodomains: The ectodomain of the SERK1 co-receptor kinase (PDB-ID 4lsc) (Santiago *et al.*, 2013) has a DALI Z-score of 21.6 and superimposes with SOBIR1 with a root mean-square deviation (r.m.s.d.) of ∼ 1 Å comparing 123 corresponding C_α_ atoms (shown in yellow in Fig. 4*a*). SERK1 shares the number of LRRs and the N-terminal capping domain with SOBIR1, but has a canonical, disulphide bond-stabilized C-terminal cap (Hohmann *et al.*, 2017). The peptide-ligand sensing PRK6 ectodomain (PDB-ID 5yah, DALI Z-score 19.2) (Zhang *et al.*, 2017) superimposes with an r.m.s.d. of ∼ 1.2 Å comparing 99 corresponding C_α_ atoms (shown in purple in Fig. 4*a*). The ectodomain of the BIR3 LRR receptor pseudokinase (PDB-ID 6fg8, DALI Z-score 20.7) aligns with an r.m.s.d. of ∼ 1.2 Å comparing 120 corresponding C_α_ atoms (shown in green in Fig. 4*a*). Together our structural comparisons reveal that functionally diverse plant receptor ectodomains share strong structural homology, with the exception of the N-terminal and C-terminal capping domains. In SOBIR1, a unique β-hairpin protrudes the N-terminal cap (Fig. 3*b*, 4*a*). This hairpin and a loop structure connecting the N-terminal capping helix with the β-hairpin both appear flexible, as judged by the analysis of the crystallographic temperature factors (Fig. 4*b*).

We located a crystallographic SOBIR1 dimer in our crystals, which would bring the C-terminal capping domains (connecting to the trans-membrane helices in the context of the full-length receptor) in close proximity (Fig. 2*a*). Analysis of the crystal packing with the PISA server (Krissinel & Henrick, 2007) revealed a rather small complex interface of ∼ 1,000 Å^2^ formed by several salt bridges, hydrogen bonds and few hydrophobic contacts. We next performed analytical size exclusion chromatography and right-angle light scattering experiments to assess the oligomeric state of the SOBIR1 extracellular domain in solution. At pH 5.0 (which corresponds to the pH associated with the plant cell wall compartment), we found our SOBIR^1-270^ construct used for crystallisation to be a monodisperse monomer with an approximate molecular weight of 30.2 kDa (calculated molecular weight is 27.4 kDa) (Fig. 5*a,b*). A longer construct that includes the complete extracellular region of SOBIR1 up to the trans-membrane helix (residue 1-283) also behaves as a monomer with an observed molecular weight of 38.1 kDa (calculated molecular weight is 33.3 kDa) (Fig. 5 *a,b*). These in solution experiments suggest that the SOBIR1 dimer observed in our structure likely represents a crystal packing artefact.

**Figure 5.**
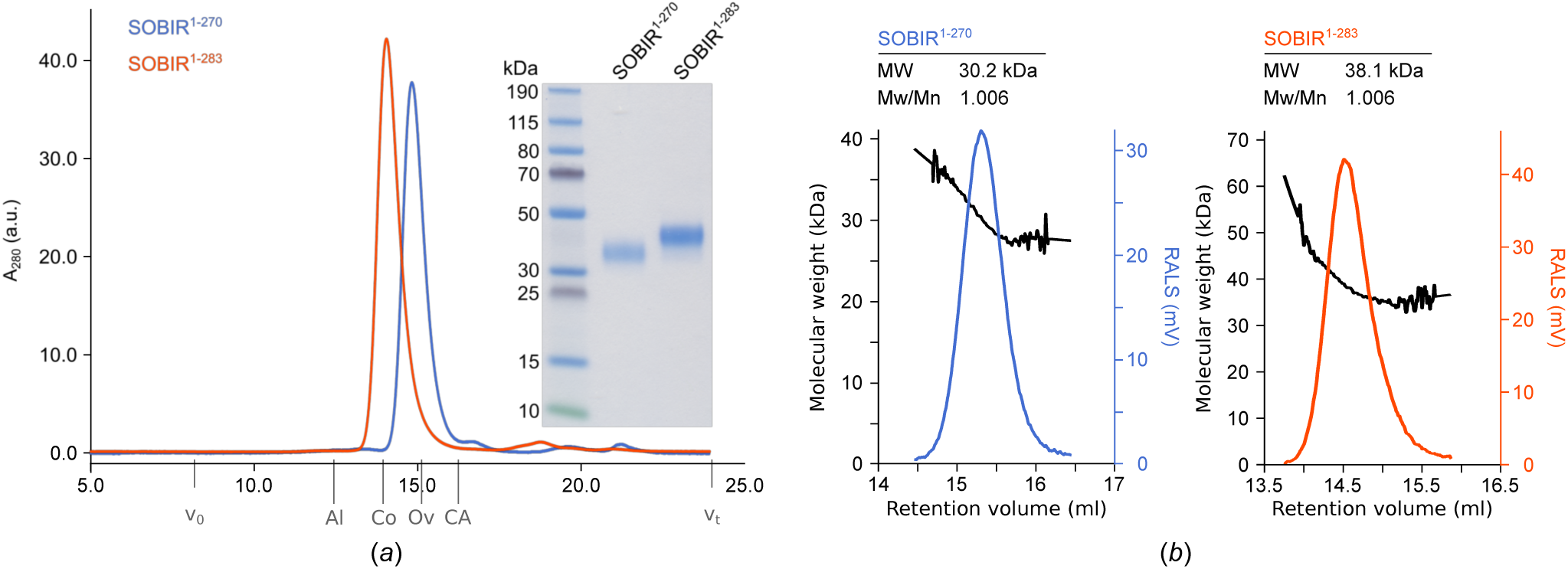
The AtSOBIR1 ectodomain is a monomer in solution. (*a*) Analytical size exclusion chromatography traces of SOBIR1^1-270^ (blue) and SOBIR1^1-283^ (orange) with the SDS-PAGE analysis of pooled peak fractions alongside. Indicated are the void volume (v_0_), the total column volume (v_t_) and the elution volumes for molecular-mass standards (Al, Aldolase, 158 kDa; Co, Conalbumin, 75 kDa; Ov, Ovalbumin, 43 kDa; CA, Carbonic Anhydrase, 29 kDa). (*b*) Analysis of the oligomeric state of AtSOBIR1. Shown are raw right-angle light scattering traces (blue and orange) and extrapolated molecular weight (black) of SOBIR1^1-270^ and SOBIR1^1-283^, with a summary table including the observed molecular weight (MW) and the dispersity (Mw/Mn) alongside. The theoretical molecular weight is 27.4 kDa for SOBIR1^1-270^ and 33.3 kDa for SOBIR1^1-283^.

**Figure 6.**
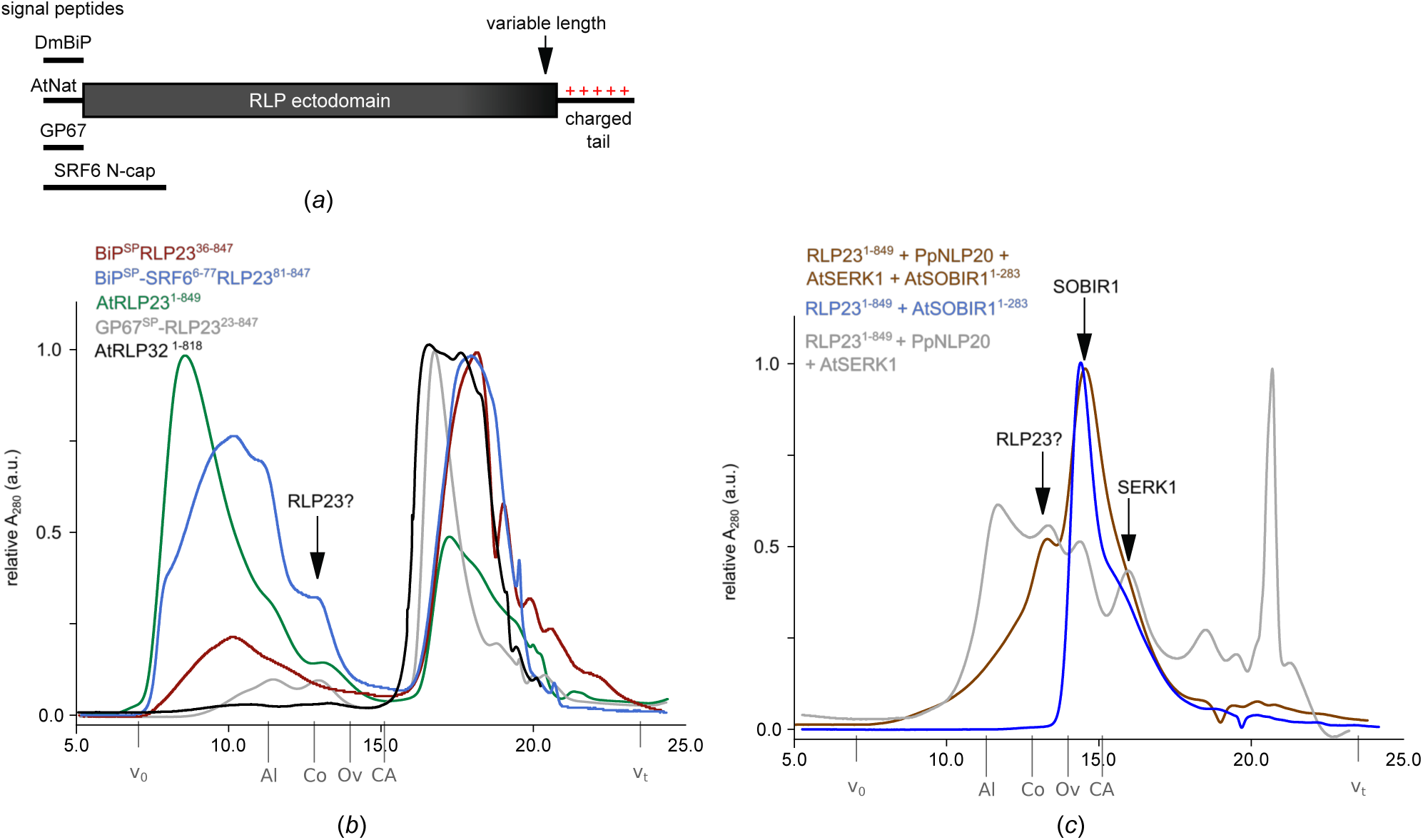
Expression and purification of AtRLPs (*a*) Schematic representation of different RLP expression constructs. Variations include different signal peptides (DmBIP, signal peptide from *Drosophila melanogaster* binding protein; AtNat, native *Arbidopsis thaliana* signal peptide, GP67, baculoviral glycoprotein 67 signal peptide, SRF6 N-cap, utilization of the whole SRF6 N-terminal capping domain) and variable construct lengths (including or omitting the charged C-terminal tail). (*b*) Example analytical size exclusion chromatography traces from RLP purifications. A peak for a monomeric, neither aggregated nor degraded RLP ectodomain (MW ∼ 100 kDa) would be expected at a elution volume of ∼ 13 ml (indicated by an arrow). Indicated are the void volume (v_0_), the total column volume (v_t_) and the elution volumes for molecular-mass standards (Al, Aldolase, 158 kDa; Co, Conalbumin, 75 kDa; Ov, Ovalbumin, 43 kDa; CA, Carbonic Anhydrase, 29 kDa). (*c*) Example analytical size exclusion chromatography traces for AtRLP23 purifications from co-expression with SOBIR^1-283^ alone (blue), with the RLP23 ligand PpNLP20 and the co-receptor AtSERK1 (grey) and with SOBIR, SERK1 and the ligand PpNLP20 (brown). Labels as in (*b*).

We next sought to test the genetic and *in vivo* biochemical finding that SOBIR1 would form heteromeric complexes with RLPs. To this end, we produced the LRR ectodomains of RLP23 and RLP32 from *Arabidopsis thaliana* by secreted expression in insect cells (see Methods). Utilisation of the native signal peptide (AtNat), the baculoviral glycoprotein 67 signal peptide (GP67), or the signal peptide from *Drosophila melanogaster* binding protein (DmBiP) (Fig. 5*a*) all lead to accumulation and secretion of RLP23, but we found the protein either aggregated or degraded in analytical size-exclusion chromatography assays (Fig. 5*b*). RLP32 showed a similar behaviour when expressed using the different signal peptides (Fig. 5*b*). We next varied the C-terminus of these constructs, omitting the positively charged C-terminal tail (Fig. 5*a*). This however did not improve the behaviour of the resulting recombinant proteins. We next replaced the flexible N-terminal capping domain of RLP23 with the N-terminal cap of the STRUBBELIG RECEPTOR FAMILY 6 LRR-RK but the resulting chimeric protein still was rapidly degraded in our preparations. Finally, we co-expressed RLP23^1-849^ with its *bona fide* peptide ligand NLP20 (Böhm *et al.*, 2014), the SOBIR1^1-283^ ectodomain, and the putative SERK co-receptor kinase (Albert *et al.*, 2015). However, also co-expression of the entire putative signalling complex did not improve the biochemical behaviour of RLP23, and we thus could not assess the role of the SOBIR1 ectodomain in immune complex formation.

## 4. Discussion

The plant membrane receptor kinase SOBIR1 is a central regulator of plant immunity. It is required for signal transduction of conserved microbe-associated molecular patterns sensed by receptor-like proteins, which lack cytoplasmic signalling domains (Gust & Felix, 2014). In genetic terms, deletion of BIR1 or over-expression of SERK3 leads to an over-activation of immune signalling (Gao *et al.*, 2009; Domínguez-Ferreras *et al.*, 2015). In both cases, these effects can be suppressed by deletion of SOBIR1, suggesting that a RLP-SOBIR-SERK complex, negatively regulated by BIR1, controls this immune response in wild-type plants. SOBIR1 and RLPs likely form heteromeric signaling complexes. Conserved GxxxG motifs in their trans-membrane regions, but neither the SOBIR1 LRR ectodomain nor the kinase domain are required for this interaction to occur *in planta* (Bi *et al.*, 2016). However, both an active kinase domain and the SOBIR1 LRR ectodomain are required for signalling (Bi *et al.*, 2016; van der Burgh *et al.*, 2019). The 1.55 Å crystal structure reveals that the SOBIR1 ectodomain has five LRRs sandwiched between unusual N-terminal and C-terminal capping domains (Fig. 2).

The known genetic *sobir1-8* missense allele (Gao *et al.*, 2009) maps to the surface of LRR2, a unique β-hairpin structure presents conserved aromatic amino acids (Phe73, Tyr85) at the surface of the domain, and the inner surface of the LRR core contains conserved patches of basic residues. Together, these structural observations argue for a role of the SOBIR1 ectodomain in mediating protein-protein interactions at the cell surface. In this respect it is of note that the area corresponding to the disordered loop region in the SOBIR1 N-terminal cap (shown in grey in Fig. 2*b*) is involved in receptor-ligand interactions in the structurally related SERK LRR-RKs (Santiago *et al.*, 2013; Sun, Li *et al.*, 2013; Santiago *et al.*, 2016; Hohmann *et al.*, 2017; Hohmann, Santiago *et al.*, 2018).

At this point we can only speculate about the nature of these protein-protein interactions. RLPs involved in plant development have been shown to directly interact with the ectodomains of large ligand-binding LRR-RKs, contributing to the sensing of small protein hormones (Lin *et al.*, 2017). We speculate that the ectodomain of SOBIR1 may play a similar role in RLP-mediated immune signalling, potentially by contributing conserved interaction surfaces from the LRR core, and/or from the protruding β-hairpin, as seen in our artificial crystallographic dimer (Figs. 2*b*, 5). In fact, the rather basic inner surface of the SOBIR1 LRR core (Fig. 3*e*) may provide a docking platform for the highly negatively charged sequence stretch in different RLPs located adjecent the RLP C-terminal capping domain (Gust & Felix, 2014).

Alternatively, the SOBIR1 ectodomain could represent a binding platform for pathogen or plant cell-wall derived ligands, based on the structural and biochemical observation that LRR domains with few repeats such as the plant RPK6 or animal lymphocyte receptors have evolved to bind peptide and small molecule ligands (Zhang *et al.*, 2017; Han *et al.*, 2008).

To test these various hypotheses, we expressed different RLPs from Arabidopsis for biochemical interaction studies, as previously reported for the RLP23 ectodomain (Albert *et al.*, 2015). However, using different expression and purification strategies (different signal peptides, construct lengths and co-expression with a secreted peptide ligand, SOBIR1 and SERKs) we could not obtain well behaving samples of RLP23 or RLP32 for quantitative binding assays. In our hands, it it thus presently not possible to dissect the contribution of the SOBIR1 ectodomain to RLP ligand sensing, complex formation and signalling at the biochemical and structural level.

## Acknowledgements

We thank T. Nürnberger and L. Zhang for providing their RLP23 expression construct for testing, and staff at beam line X06DA (PXIII) of the Swiss Light Source (SLS) Villigen, Switzerland for technical help during data collection. This work was supported by the ‘SICOPID’ ERA-CAPS network fund (to M.H., T. Nürnberger, C. Zipfel, Z. Nimchuk & Y. Jaillais) with Swiss National Science Foundation grant no. 31CP30_180213). The crystallographic coordinates and structure factors have been deposited with the Protein Data Bank (http://rcsb.org) ID 6r1h. Native and sulphur SAD diffraction images and XDS processing files have been deposited at zenodo.org with doi:10.5281/zenodo.2594485 and doi:10.5281/zenodo.2595891, respectively.

